# Epistasis lowers the genetic barrier to SARS-CoV-2 neutralizing antibody escape

**DOI:** 10.1101/2022.08.17.504313

**Authors:** Leander Witte, Viren Baharani, Fabian Schmidt, Zijun Wang, Alice Cho, Raphael Raspe, Maria C. Guzman-Cardozo, Frauke Muecksch, Christian Gaebler, Marina Caskey, Michel C. Nussenzweig, Theodora Hatziioannou, Paul D. Bieniasz

**Author notes:** Equal contribution (Leander Witte, Viren Baharani, Fabian Schmidt).

## Abstract

Consecutive waves of SARS-CoV-2 infection have been driven in part by the repeated emergence of variants with mutations that confer resistance to neutralizing antibodies Nevertheless, prolonged or repeated antigen exposure generates diverse memory B-cells that can produce affinity matured receptor binding domain (RBD)-specific antibodies that likely contribute to ongoing protection against severe disease. To determine how SARS-CoV-2 omicron variants might escape these broadly neutralizing antibodies, we subjected chimeric viruses encoding spike proteins from ancestral, BA.1 or BA.2 variants to selection pressure by a collection of 40 broadly neutralizing antibodies from individuals with various SARS-CoV-2 antigen exposures. Notably, pre-existing substitutions in the BA.1 and BA.2 spikes facilitated acquisition of resistance to many broadly neutralizing antibodies. Specifically, selection experiments identified numerous RBD substitutions that did not confer resistance to broadly neutralizing antibodies in the context of the ancestral Wuhan-Hu-1 spike sequence, but did so in the context of BA.1 and BA.2. A subset of these substitutions corresponds to those that have appeared in several BA.2 daughter lineages that have recently emerged, such as BA.5. By including as few as 2 or 3 of these additional changes in the context of BA.5, we generated spike proteins that were resistant to nearly all of the 40 broadly neutralizing antibodies and were poorly neutralized by plasma from most individuals. The emergence of omicron variants has therefore not only allowed SARS-CoV-2 escape from previously elicited neutralizing antibodies but also lowered the genetic barrier to the acquisition of resistance to the subset of antibodies that remained effective against early omicron variants.

## Introduction

Consecutive waves of SARS-CoV-2 infection have been driven in part by the repeated emergence of variants with mutations that confer resistance to neutralizing antibodies[1-8] Nevertheless, prolonged or repeated antigen exposure generates diverse memory B-cells [9-14] that can produce affinity matured receptor binding domain (RBD)-specific antibodies that likely contribute to ongoing protection against severe disease. RBD-specifiic antibodies are the dominant source of neutralizing activity in the plasma of infected or vaccinated individuals[15, 16] and can be broadly grouped into 4 prototype classes[17, 18]. Class 1 and 2 antibodies bind epitopes overlapping the ACE-2 binding site (Fig. 1a), while class 3 and 4 antibodies bind outside the ACE-2-binding site on opposite sides of the RBD. The emergence of viral variants, particularly omicron BA.1 and BA.2 with numerous RBD substitutions has reduced the effectiveness of neutralizing antibodies as components of the convalescent and vaccine-elicited immune response and as therapeutics[3-8]. However, the retention of activity against emergent variants, such as BA.1 and BA.2 and derivatives thereof [4, 14, 19, 20], by a subset of plasma and memory antibodies likely contributes to the residual effectiveness of ancestral variant-based vaccines, and the partial protection afforded by exposure to antigens from earlier variants. A key question is therefore is how SARS-CoV-2 might evolve to evade these diverse and broadly neutralizing antibodies.

**Figure 1:**
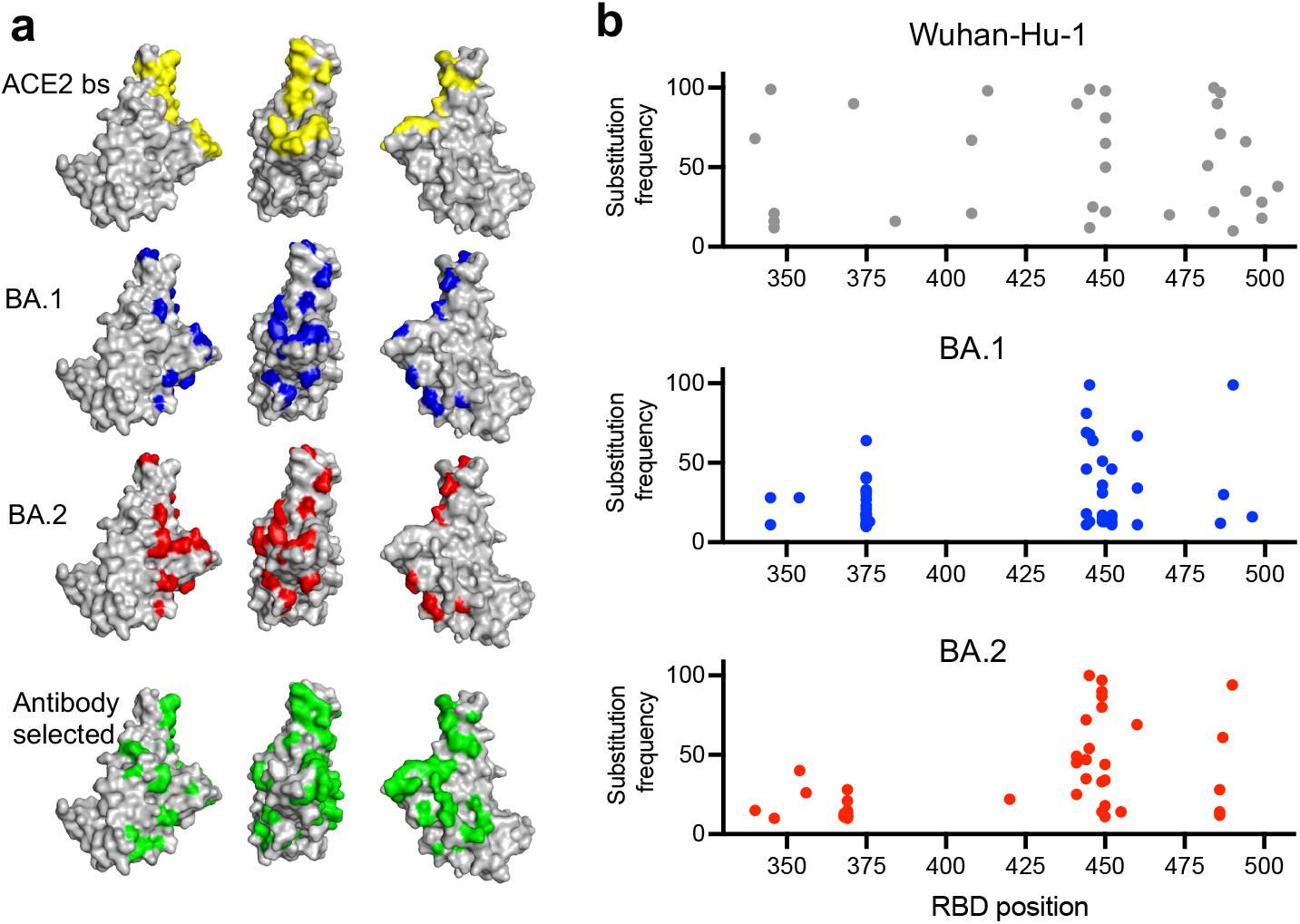
Selection of SARS-CoV-2 spike variants with antibodies. **a)** RBD structure (PDB ID : 7C8J) illustrating salient features: yellow = ACE2 binding site; blue = residues changed in BA.1 relative to Wuhan-Hu-1; red = residues changed in BA.2. Highlighted in green are all positions where substitutions were apparently selected by neutralizing antibodies during replication of rVSV/SARS-CoV-2 encoding Wuhan-Hu-1, BA.1, or BA.2 spike proteins. **b)** Substitution frequency at residues along the length of the RBD following passage of rVSV/SARS-CoV-2 encoding Wuhan-Hu-1, BA.1, and BA.2 spike proteins with antibodies.

### Selection of SARS-CoV-2 spike variants with antibodies

We chose 40 antibodies, isolated from the memory B-cells of multiple volunteer cohorts with varying exposures to SARS-CoV-2 antigens, using an ancestral Wuhan-Hu-1 RBD (Supplementary Data 1 and 2). Specifically, the antibodies were from (i) participants infected with Wuhan-Hu-1 at 6.2 or 12 months previously, some of whom had also received 1 or 2 mRNA vaccine doses prior to antibody isolation[9, 13] (ii) recipients of 3 mRNA vaccine doses who had not been infected (antibodies isolated at ∼1m after the third dose)[14] or (iii) individuals who had experienced an omicron BA.1 breakthrough infection after 3 vaccine doses (antibodies isolated at ∼1m after infection)[21, 22]. We chose ‘broadly’ neutralizing antibodies, defined as those that could neutralize both Wuhan-Hu-1 and omicron (BA.1) pseudotyped viruses, with IC_50_ values of ∼100ng/ml or less. Most of these antibodies (35 of 40) also neutralized BA.2. With the exception of two class 1/4 broadly neutralizing antibodies, isolated from infected individuals at 1.3m post-infection[18, 23], the antibodies were chosen without regard to any property (e.g. V_H_,V_L_ or RBD epitope class, Supplementary Data 2) other than potency and breadth. Based on competition experiments with prototype class 1 through 4 antibodies, the broadly neutralizing collection targeted diverse RBD epitopes that included all classes, but exhibited some skew toward class 3.

To select antibody escape mutants, we employed replication competent recombinant vesicular stomatitis viruses (rVSV/SARS-CoV-2)[24] that encoded the spike proteins from ancestral Wuhan-Hu-1, or BA.1 or BA.2 variants in place of G. Diversified rVSV/SARS-CoV-2 populations containing 10^6^ infectious units were incubated with antibodies at 1 µg/ml before infection of target cells. Following 2 passages in the presence of each antibody, the acquisition of RBD substitutions in the selected progeny viral populations was evaluated by next-generation sequencing (Supplementary Data 3). There was no apparent difference in the ability of antibodies from different classes to select mutations. Indeed, in a total of 120 selection experiments, 39 out of the 40 antibodies tested yielded substitutions, present at a frequency of >10% of sequences in the selected viral populations, Overall, these substitutions were at 34 different positions in the RBD (Fig. 1a, b, Supplementary Fig. 1). Substituted positions in the selected viral populations were spatially proximal to, or within, the ACE2 binding site and were largely consistent with the selecting antibody class designation, determined by competition with prototype class 1-4 antibodies (Supplementary Data.2, Supplementary Fig. 1). With only 3 exceptions, two of which were reversion of BA.1 specific substitutions (F375S and S496G), the antibody selection experiments in the BA.1 and BA.2 context generated substitutions at positions that were not already changed in BA.1 or BA.2 compared to Wuhan-Hu-1 (Fig. 1a, b, Supplementary Figure 1).

### Broadly neutralizing antibody resistance mutations

We chose 23 substitutions that were enriched during passage of rVSV/SARS-CoV-2 variants in the presence of the 40 antibodies and constructed spike plasmids based on the context in which the substitutions were selected. If a substitution was enriched in the ancestral rVSV/SARS-CoV-2_Wuhan-Hu-1_ context, a corresponding mutant Wuhan-Hu-1 spike expression plasmid was constructed. In instances where a position was substituted in either of the antibody selected rVSV/SARS-CoV-2_BA.1_ or rVSV/SARS-CoV-2_BA.2_ populations the corresponding position was substituted in all three parental spike expression plasmids. Thus, we generated 56 pseudotyped HIV-1 viruses with mutant spike sequences and tested their ability to resist neutralization by each of the 40 broadly neutralizing antibodies, at a concentration of 1 µg/ml (Fig. 2a, Supplementary Data 4). Infection was quantified relative to uninhibited virus (absence of antibody), and antibody ‘escape’ was defined as a 5-fold increase in infection relative to the antibody-inhibited parental pseudotype, provided infection reached >10% of the uninhibited control. Most of the substitutions conferred specific resistance to one or more antibodies of the same class, consistent with their position on the RBD surface. For example, class 4 or 1/4 broadly neutralizing antibodies, which bind epitopes that are well conserved in sarbecoviruses[18] were escaped by mutations at P384, R408, G413, or G504 (Fig 2a, Supplementary Fig.2). Notably, some class 1/4 or 4 broadly neutralizing antibodies were ineffective or less potent against BA.2, which has substitutions that encroach on class 4 epitopes (Supplementary Fig.2). While most substitutions resulted in class-specific antibody resistance, two substitutions positioned at the base of the RBD either gave modest class-independent reductions in sensitivity (F375S), or no reductions in antibody sensitivity (Y369F). We speculate that these represent fitness-enhancing substitutions that arose during the rVSV/SARS-CoV-2 passage.

**Figure 2:**
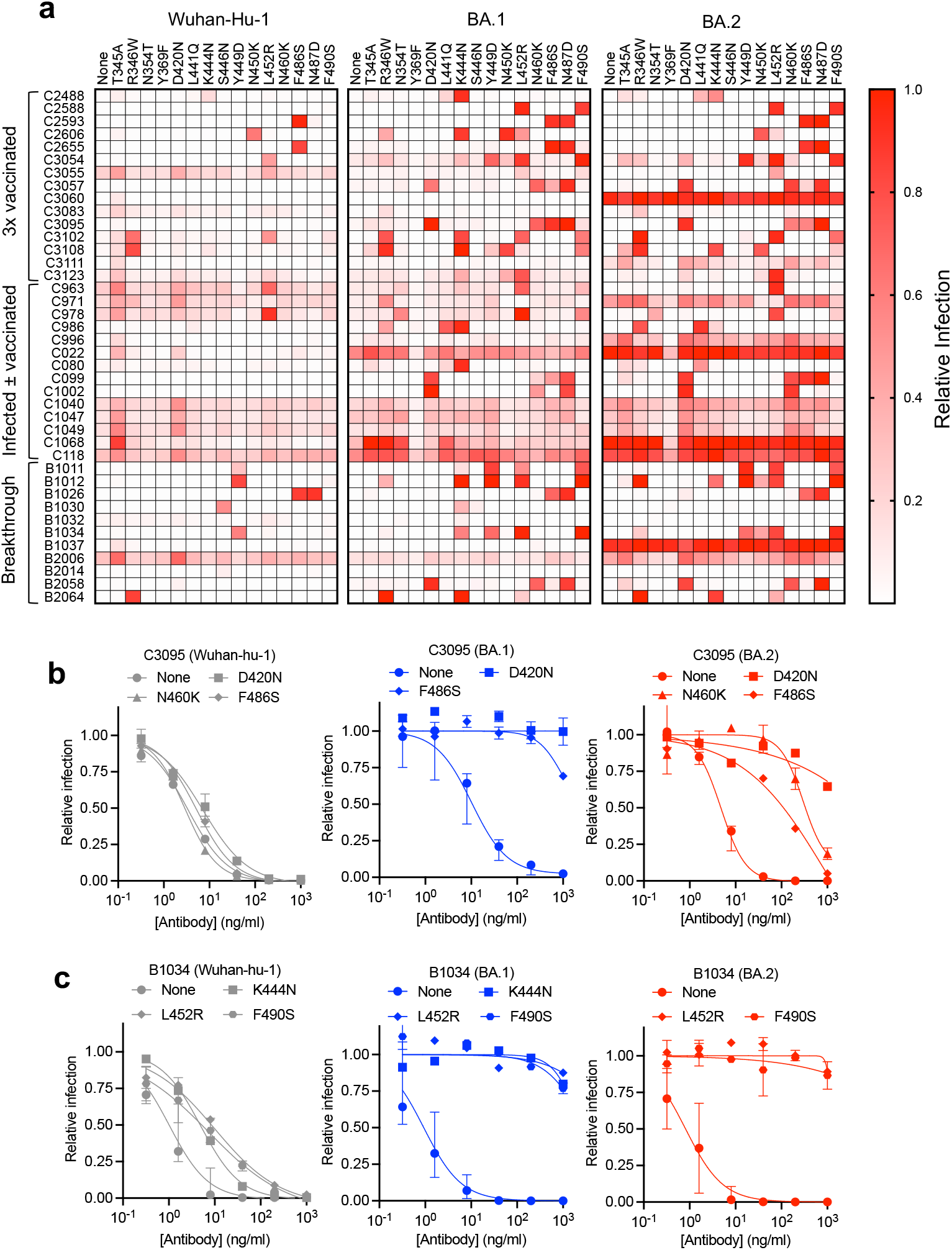
Epistasis and neutralizing antibody resistance. **a)** Inhibition of Wuhan-Hu-1, BA.1, and BA.2 RBD point mutant pseudotypes by broadly neutralizing antibodies. Relative infection is defined as the decimal fraction of infection measured (with 1 μg/ml antibody), relative to an uninhibited virus control (without antibody). Median value of two experiments. **b)** Neutralization of the RBD point mutant pseudotypes in Wuhan-Hu-1, BA.1, and BA.2 backgrounds by antibody C3095. **c)** Neutralization of the RBD point mutant pseudotypes indicated in Wuhan-Hu-1, BA.1, and BA.2 backgrounds by antibody B1034. For **b** and **c** the median value and range of two or three independent experiments is shown.

### Epistasis and neutralizing antibody resistance

For more than half of the broadly neutralizing antibodies (21 out of 40), we found single-amino-acid substitutions that conferred antibody resistance in the context of BA.1 or BA.2 pseudotypes, but not in the context of Wuhan-Hu-1 pseudotypes. Thus, substitutions introduced in the BA.1 and BA.2 contexts led to escape from a greater number of antibodies than the same substitutions in the Wuhan-Hu-1 context (Fig. 2a, Supplementary Fig. 3). For many of these substitutions, we confirmed their effects by determining full neutralization curves using Wuhan-Hu-1, BA.1 and BA.2 pseudotypes (Fig. 2b, c, Supplementary Fig. 4, 5, 6). For example, substitutions D420N, N460K and F486S caused negligible changes in sensitivity to the class 1/2 antibody C3095 in the Wuhan-Hu-1 context but conferred complete resistance or large potency deficits (10 to 100-fold increased IC_50_) in the BA.1 or BA.2 context (Fig. 2b). Similarly, for the class 3 antibody B1034, the K444N, L452R and F490S substitutions conferred modest potency deficits for Wuhan-Hu-1 pseudotypes, but near-complete loss of neutralization in the BA.1 and/or BA.2 contexts (Fig. 2c).

Similar epistatic effects were evident for 14 of the 15 antibody resistance substitutions that were tested in all three pseudotype backgrounds (Fig 2a, Supplementary Fig. 4, 5, 6, Supplementary Data 4). Substitutions at positions N354, F375, D420, L441, N460 and F490 conferred resistance to one or more antibodies when introduced into BA.1 or BA.2 but did not do so when introduced into Wuhan-Hu-1. For 11 antibodies, single amino acid substitutions (at positions T345, R346, K444, Y449, N450, L452, F486 or N487) conferred resistance in a context-independent manner. However, the very same substitutions conferred resistance to 10 other antibodies when introduced into BA.1 and/or BA.2 but not when introduced into the Wuhan-Hu-1. Overall, there was extensive epistatic interaction between antibody resistance substitutions and variation in the BA.1 or BA.2 proteins (Fig. 2a, Supplementary Data 4).

Included in the set of broadly neutralizing antibodies was C099 (class 1), for which a structure of the Fab:RBD interface has been determined[12], and for which we failed to generate single amino-acid resistance mutations in the context of Wuhan-Hu-1. In contrast, C099 resistance was readily generated by single amino acid substitutions in the BA.1 and BA.2 contexts (Fig 2a, Supplementary Data 4). Indeed, two substitutions (at D420 and N460) within or proximal to the target epitope (Supplementary Fig. 7a) were necessary to confer resistance to C099 in the Wuhan-Hu-1 context[12], but each of these substitutions, as well as F486S or N487D, could individually lead to full C099 resistance in BA.1 and/or BA.2 (Fig 2a, Supplementary Fig. 4, Fig. 7a). Similarly, single target epitope substitutions at positions R346, K444 and L441 conferred resistance to C032, a clonal ancestor of the affinity-matured class 3 broadly neutralizing antibody, C080 (Supplementary Fig. 7b) [12], but none conferred resistance to C080 in the context of Wuhan-Hu-1 (Fig 2a, Supplementary Fig 5). Again, however, these single substitutions could generate C080 resistance in the BA.1 or BA.2 background. Overall, we conclude that BA.1 and BA.2 variants more readily escape broadly neutralizing antibodies such as C099 and C080 because they already possess a subset of the multiple target epitope substitutions required for antibody resistance (Supplementary Fig. 7a, b).

### Mutations selected *in vitro* and BA.2 daughter lineages

Recently, sublineages derived from BA.2 have displaced BA.1 and BA.2[25, 26]. These sublineages have thus far exhibited varying degrees of prevalence, but most encode RBD substitutions, including L452Q (BA.2.12), G446S, N460K (BA.2.75), L452R, F486V (BA.4 and BA.5) and R346T (BA.4.6)[25, 26]. These substitutions are identical to, or at the same positions as, substitutions exhibiting epistatic interactions to confer resistance to multiple broadly neutralizing antibodies (Fig 2a, Supplementary Fig 4). In particular, we found that substitutions at positions that changed during the BA.2 to BA.5 transition conferred resistance to antibodies of multiple classes (class 2/3 and 3 for L452R and classes 1, 1/2 and1/4 for F486S). Notably, for 7 of 14 antibodies, resistance conferred by L452 or F486 substitutions was dependent on prior acquisition of substitutions in BA.2 (Supplementary Fig. 8).

### Resistance of synthetic SARS-CoV-2 variants to broadly neutralizing antibodies

Synthetic SARS-CoV-2 ‘polymutant’ spike proteins, encoding combinations of changes arising during *in vitro* selections experiments, can have emergent escape characteristics similar to those of natural SARS-CoV-2 variants[4, 16]. We generated synthetic RBD variants with combinations of mutations selected *in vitro* to assess the potential effect of selective pressure by broadly neutralizing antibodies. Specifically, we compiled RBD mutations that conferred resistance to non-overlapping subsets of antibodies (Fig. 2a). For instance, the R346W substitution conferred resistance to several class 3 antibodies (e.g. C3102, C3108, C1068), the D420N substitution enabled escape from several class 1/4 antibodies (e.g. C099, C1002, C3095, C3111), while the K444N substitution enabled escape from various antibodies particularly in class 2/3 (e.g. B1011, B1012, B1034). In some cases, two or more alternative substitutions (e.g. T345A and R346W) conferred resistance to similar sets of antibodies (Fig. 2a). In these instances, only one substitution (R346W) was chosen. We tested seven synthetic RBD variants, with 2 to 8 substitutions introduced in the context of a BA.5 spike (BA5(+2) through BA5(+8), Fig. 3a) for neutralization by the antibody panel. The synthetic variants all generated infectious pseudotypes, and all resisted neutralization by 32 or more of the 40 antibodies, across all classes (Fig. 3b, c). Notably, the BA.5(+2) and BA.5(+3) variants, with only two (R346W, D420N) and three (R346W, D420N, K444N) substitutions, were resistant to 36 and 38 of the 40 broadly neutralizing antibodies, respectively. Overall, the 40 broadly neutralizing antibodies inhibited Wuhan-Hu-1, BA.1 and BA.2 pseudotype infection by a median of 170-, 40- and 118-fold at 1μg/ml, respectively (Fig. 3c). For BA.5, the median inhibition was 6-fold, while for each of the synthetic variants, median inhibition was only 1.0- to 1.2-fold. We conclude that compared to early omicron variants, few of the broadly neutralizing RBD antibodies examined herein remain effective against potential future SARS-CoV-2 variants derived from BA.5 with a limited number of additional substitutions.

**Figure 3.**
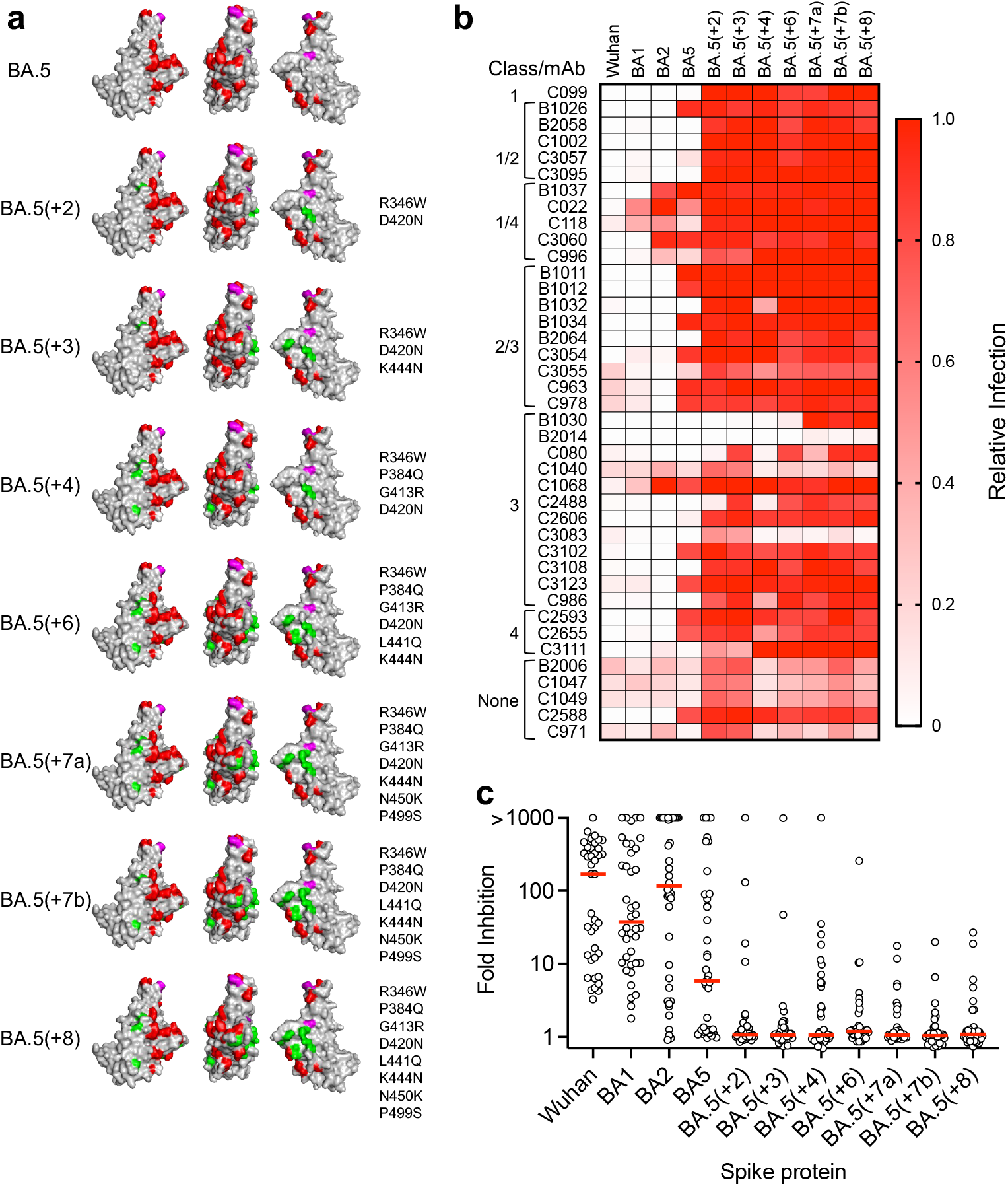
Resistance of synthetic SARS-CoV-2 variants to broadly neutralizing antibodies. **a)** BA.5 and synthetic SARS-CoV-2 variants. Red: Substitutions in BA.2 relative to Wuhan-Hu-1; magenta: substitutions BA.5 relative to BA2. Highlighted in green are substitutions in the synthetic variants, chosen based on antibody selection experiments. **b)** Inhibition of synthetic variants by broadly neutralizing antibodies. Relative infectivity was calculated as a decimal fraction of infection measured (with 1 µg/ml antibody) relative to an uninhibited virus control (without antibody). A mean of two independent experiments is shown. **c)** Fold inhibition of natural and synthetic variant infection by individual broadly neutralizing antibodies. Individual data points represent the median of two independent experiments for individual antibodies. Red line = group median fold-inhibition for all antibodies tested.

### Plasma neutralization of synthetic SARS-CoV-2 variants

Populations of antibodies in the memory B-cell compartment may differ from those represented in plasma[27]. We selected 3 synthetic variants (BA.5(+3), BA.5(+7b) and BA.5(+8)) and determined their neutralization sensitivity to plasma antibodies from individuals with heterogeneous antigen exposures (Fig 4, Supplementary Data 1, 5). Individuals who were infected with Wuhan-Hu-1-like viruses (but not vaccinated) exhibited low plasma NT_50_ against all omicron variants, as did those whose only antigen exposure was 2 doses of an mRNA vaccine. These groups had median NT_50_ titers of 96-257 for BA.1 and BA.2 variants and close to background NT_50_ values for BA.5 and the synthetic BA.5-based variants (Fig. 4).

**Figure 4:**
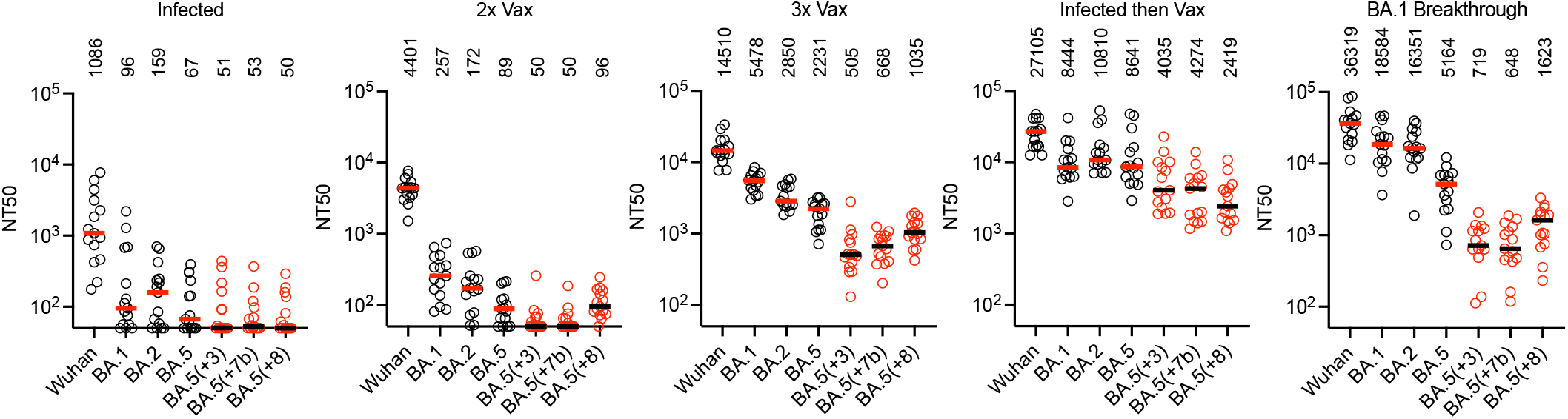
Reduced potency of polyclonal plasma antibodies against synthetic variants. The 50% neutralization titers (NT_50_) for plasmas from various volunteer cohorts (see methods). Fifteen randomly selected plasma samples from each cohort were tested against natural variants; Wuhan-Hu-1, BA.1, BA.2, BA.5 (black), and synthetic variants; BA.5(+3), BA.5(+7b) BA.5(+8) (red). Individual points represent individual plasma samples; median of two independent experiments is plotted. Group median NT_50_ is indicated by red/black lines and numbers above each group.

Plasma from three groups who had received multiple antigen exposures, (i) 3x vaccinated, (ii) infected then vaccinated or (ii) 3x vaccinated and breakthrough infection – the same groups from which the broadly neutralizing antibodies were obtained (Supplementary Data 1) – had higher neutralizing titers against BA.1, BA.2 and BA.5 (NT_50_ values of 2231-18584). The synthetic variants were less sensitive to neutralization by these plasmas than either of the BA.2 or BA.5 parental variants (Fig. 4). For example, plasmas from 3x vaccinated individuals, had median NT_50_ values against BA.5(+3) that were 6-fold and 4-fold reduced compared to BA.2 and BA.5 respectively. Notably, the group with BA.1 breakthrough infection following 3x vaccination had higher NT_50_ values against BA.1 (3.4-fold) and BA.2 (5.7-fold) than those without breakthrough infection (Fig. 4). However, these apparent gains in neutralizing activity after BA.1 breakthrough infection were absent against the synthetic variants.

Consequently, the BA.5(+3) variant was 23-fold and 7-fold less sensitive than BA.2 and BA.5 to plasmas from the BA.1 breakthrough infection group (Fig. 4). We conclude that a significant portion of the residual BA.5 plasma neutralizing activity in these groups can be evaded by the acquisition of a few additional substitutions.

Notably, plasma from individuals who had been infected early in the pandemic and subsequently vaccinated (and were thus exposed to ancestral Wuhan-Hu-1-like antigens)[13] had broader neutralization properties than the 3x vaccinated and BA.1 breakthrough groups (Fig. 4). Indeed, for the infected then vaccinated group, the decrease in median NT_50_ for BA.5 compared to BA.2 was not statistically significant, and the synthetic variants were only 2-fold to 3.6-fold less sensitive than BA.5 to these plasmas. These decrements reached significance only for the BA.5(+7b) and BA.5(+8) variants (p-values 0.02 and 0.01, respectively, Supplementary Data 5).

## Discussion

Prolonged or repeated exposure to antigen drives somatic mutation of antibody lineages resulting in greater diversity and affinity and, in the context of SARS-CoV-2, greater neutralization breadth[9-14]. A key consequence of antibody maturation is that SARS-CoV-2 escape from neutralization by individual RBD antibodies often requires more than one substitution in the target epitope[12]. A corollary, described herein, is that epistatic interactions between newly acquired and pre-existing substitutions can enable escape from neutralizing antibodies that tolerate single but not multiple target epitope substitutions. This feature is evident in BA.1 and BA.2 that encode many mutations in the class 1-4 RBD epitopes, which provide a background whereby a small number of additional RBD substitutions can result in epitope loss for most broadly neutralizing antibodies. The rapid sequential displacement of early omicron variants by subsequent derivatives may be because individual substitutions have greater impact on antibody escape in omicron than in ancestral contexts.

Analysis of antibodies in plasma and in the Wuhan-Hu-1-selected memory B-cell compartment suggest that BA.1 breakthrough infection selectively boosts the subset of vaccine elicited neutralizing antibodies residing in memory that neutralize BA.1[21, 22, 28, 29]. While some of these memory antibodies also neutralized BA.5, most did not neutralize synthetic variants with only 3 additional substitutions. BA.1 infection after 3 vaccine doses thus provides no increase in plasma neutralization over 3 vaccine doses alone against such variants. In contrast, boosting of a diverse set of evolved antibodies, elicited by ancestral Wuhan-Hu-1 infection, using a Wuhan-Hu-1-based vaccine increases, and further matures, a broader set of antibodies[13] subsets of which may cross-react with distinct variants. Our findings suggest that differences in antigen exposure may generate significant heterogeneity in antibody neutralizing breadth and potency that is increasingly evident as the antigenic distance between currently circulating and ancestral SARS-CoV-2 variants grows. Epistatic interaction between pre-existing and new viral mutations, along with population heterogeneity in neutralizing antibody responses, will likely continue to impact the future course and pace of SARS-CoV-2 variant emergence.

## Acknowlegements

We thank Anna Gazumyan and Brianna Johnson for assistance with the expression of antibodies. This work was supported by National Institutes of Health (NIH) grant P01AI165075 to PDB, TH and MCN, R37AI64003 to PDB, R01AI78788 to TH, P01-AI138398-S1 and 2U19AI111825 to MCN. FM was supported by the Bulgari Women & Science Fellowship in COVID-19 Research. PDB and MCN are Howard Hughes Medical Institute (HHMI) Investigators. This article is subject to HHMI’s Open Access to Publications policy. HHMI lab heads have previously granted a nonexclusive CC BY 4.0 license to the public and a sublicensable license to HHMI in their research articles. Pursuant to those licenses, the author-accepted manuscript of this article can be made freely available under a CC BY 4.0 license immediately upon publication.

## Methods

### Cell Culture

All cell lines were cultured in DMEM supplemented with 10% fetal calf serum and 10 µg/ml gentamicin at 37ºC and 5% CO_2_. Cells were periodically checked for mycoplasma and retrovirus contamination by DAPI staining and reverse transcriptase assays, respectively. The cell lines used have been previously described[24].

### SARS-CoV-2 variant spike expression plasmids

Generation of an omicron BA.1 spike expression plasmid lacking the C-terminal 19 amino acids and encoding the R683G furin cleavage site substitution (pCR3.1-BA.1Δ19) has been previously described[4]. Plasmids encoding similar spike proteins from the BA1.1, BA.2 and BA.4/5 variants were derived from this plasmid by an overlap extension PCR strategy followed by Gibson assembly in the pCR3.1 backbone that was linearized by NheI/XbaI digestion. The variant-specific changes introduced were:

Omicron BA.1: A67V, Δ69-70, T95I, G142D, Δ143-145, Δ211, L212I, ins214EPE, G339D, S371L, S373P, S375F, K417N, N440K, G446S, S477N, T478K, E484A, Q493K, G496S, Q498R, N501Y, Y505H, T547K, D614G, H655Y, H679K, P681H, N764K, D796Y, N856K, Q954H, N969H, N969K, L981F Omicron BA.1.1: A67V, Δ69-70, T95I, G142D, Δ143-145, Δ211, L212I, ins214EPE, G339D, R346K, S371L, S373P, S375F, K417N, N440K, G446S, S477N, T478K, E484A, Q493K, G496S, Q498R, N501Y, Y505H, T547K, D614G, H655Y, H679K, P681H, N764K, D796Y, N856K, Q954H, N969H, N969K, L981F Omicron BA.2: T19I, L24S, del25-27, G142D, V213G, G339D, S371F, S373P, S375F, T376A, D405N, R408S, K417N, N440K, S477N, T478K, E484A, Q493R, Q498R, N501Y, Y505H, D614G, H655Y, N679K, P681H, N764K, D796Y, Q954H, N969K Omicron BA.5: T19I, L24S, del25-27, del69-70, G142D, V213G, G339D, S371F, S373P, S375F, T376A, D405N, R408S, K417N, N440K, L452R, S477N, T478K, E484A, F486V, Q498R, N501Y, Y505H, D614G, H655Y, N679K, P681H, N764K, D796Y, Q954H, N969K

### Monoclonal antibodies and plasma samples

Monoclonal antibodies were cloned from the memory B-cells of individuals with varying exposure to SARS-CoV-2 antigen as a result of infection and/or vaccination.

Plasma samples were from 5 groups

(i) Individuals who were infected with Wuhan-Hu-1-like viruses ∼1y prior to blood collection, but not vaccinated[13].

(ii) Individuals who received 2 doses of mRNA vaccine (2^nd^ dose ∼1m prior to plasma collection) [30]

(iii) Individuals who received 3 doses of mRNA vaccine (3^rd^ dose ∼1m prior to plasma collection) [14]

(iii) individuals from the same Wuhan-Hu-1-like infection cohort who were infected ∼1y previously and also received 2 mRNA vaccine doses prior to blood collection[13].

(iii) individuals who received 3 mRNA vaccine doses and were then infected by omicron BA.1 ∼ 1m prior to blood collection [21, 22].

The study visits and blood draws were obtained with informed consent from all participants under a protocol that was reviewed and approved by the Institutional Review Board of the Rockefeller University (IRB no. DRO-1006, ‘Peripheral Blood of Coronavirus Survivors to Identify Virus-Neutralizing Antibodies’).

### rVSV/SARS-CoV-2/GFP construction and rescue

The generation of rVSV/SARS-CoV-2/GFP chimeric viruses encoding SARS-2 spike proteins has been previously described[24]. A plaque-purified variant designated rVSV/SARS-CoV-2/GFP_2E1_ encoding D215G/R683G substitutions was used in this study. To introduce the spike proteins of VOCs BA.1.1 and BA.2, the respective spike sequences were amplified from the pCR3.1 constructs using PCR and primers specific for the VSV backbone. The rVSV backbone was digested with MluI/XhoI and the SARS-CoV-2 spike insert was introduced by Gibson assembly.

To rescue the recombinant viruses, HEK-293T/ACE2cl.22 cells were seeded 1×10^6^/well in poly-D-lysine-coated 6-well plates one day prior to transfection. The following day, cells were rinsed with serum-free DMEM and infected with a recombinant vaccinia virus expressing T7 polymerase at an MOI of 5 for 45 min at 37ºC, gently rocking every 10-15 min. The inoculum was removed and 1.5 ml DMEM supplemented with 10% FCS was added. Next, cells were transfected with an plasmid mixture containing the rVSV/SARS-CoV-2 genome (500 ng) and the helper plasmids pBS-N (300 ng), pBS-P (500 ng), pBS-L (100ng) and pBS-G (800 ng) using the lipofectamine LTX (9 μl) and PLUS (5.5 μl) transfection reagents. The DNA and PLUS reagent were mixed in 100 μl OptiMEM to which LTX was added in 105 μl OptiMEM. The reaction was incubated for 20 min at RT and then added to the cells. The transfected cells were monitored by microscopy and the supernatant of cells expressing GFP was collected ∼48 h post transfection, filtered using a 0.1 µm filter to remove residual vaccinia virus and transferred to HEK-293T/ACE2cl.22 cells in 6-well plates to amplify the rescued virus. After ∼48 h the supernatant of the infected cells was collected and filtered using a 0.22 µm filter and transferred onto HEK-293T/ACE2cl.22 cells seeded in T175 flasks. The viruses were passaged several times in T175 flasks to create diversity in the viral population and generate virus stocks of >10^8^ IU/ml. The titer of the viral stocks was assessed by serial dilution, infection of HEK-293T/ACE2cl.22 cells cells and subsequent flow cytometry.

### Antibody sequencing, cloning, and expression

Antibodies were identified and sequenced as previously described[23]. In brief, RNA from single cells was reverse-transcribed (SuperScript III Reverse Transcriptase, Invitrogen, 18080–044) and the cDNA stored at −20 °C or used for subsequent amplification of the variable IGH, IGL and IGK genes by nested PCR and Sanger sequencing. Sequence analysis was performed using MacVector. Amplicons from the first PCR reaction were used as templates for sequence- and ligation-independent cloning into antibody expression vectors. Recombinant monoclonal antibodies were produced and purified as previously described[23].

### Biolayer interferometry

Biolayer interferometry assays were performed as previously described[23]. In brief, we used the Octet Red instrument (ForteBio) at 30 °C with shaking at 1,000 r.p.m. Epitope binding assays were performed with protein A biosensor (ForteBio 18-5010), following the manufacturer’s protocol “classical sandwich assay” as follows: (1) Sensor check: sensors immersed 30 sec in buffer alone (buffer ForteBio 18-1105), (2) Capture 1st Ab: sensors immersed 10 min with Ab1 at 10 μg/mL, (3) Baseline: sensors immersed 30 sec in buffer alone, (4) Blocking: sensors immersed 5 min with IgG isotype control at 10 μg/mL. (5) Baseline: sensors immersed 30 sec in buffer alone, (6) Antigen association: sensors immersed 5 min with RBD at 10 μg/mL. (7) Baseline: sensors immersed 30 sec in buffer alone. (8) Association Ab2: sensors immersed 5 min with Ab2 at 10 μg/mL. Curve fitting was performed using the Fortebio Octet Data analysis software (ForteBio). Affinity measurement of anti-SARS-CoV-2 IgGs binding were corrected by subtracting the signal obtained from traces performed with IgGs in the absence of WT RBD. The kinetic analysis using protein A biosensor (as above) was performed as follows: (1) baseline: 60sec immersion in buffer. (2) loading: 200sec immersion in a solution with IgGs 10 μg/ml. (3) baseline: 200sec immersion in buffer. (4) Association: 300sec immersion in solution with WT RBD at 20, 10 or 5 μg/ml (5) dissociation: 600sec immersion in buffer. Curve fitting was performed using a fast 1:1 binding model and the Data analysis software (ForteBio). Mean KD values were determined by averaging all binding curves that matched the theoretical fit with an R2 value ≥ 0.8. To classify antibodies, competition with antibodies C105 (class 1) C144 (class 2) C135 (class 3) and CR3022 (class 4)[17, 23] was determined.

### Selection of antibody resistant rVSV/SARS-CoV-2 variants

To identify antibody escape mutations in rVSV/SARS-CoV-2/GFP_2E1_, rVSV/SARS-CoV-2/GFP_BA1.1_, and rVSV/SARS-CoV-2/GFP_BA2_, viral populations containing 1×10^6^ infectious units were incubated with monoclonal antibodies (mAbs) at a concentration of 1 µg/mL for 1 hour at 37^°^C. The virus/antibody mixture was then used to inoculate HEK-293T/ACE2cl.22 cells in 6 well plates. The next day, the medium was replaced with fresh medium containing the monoclonal antibody at 1 µg/ml. After a further 24 hours, the virus-containing supernatant was harvested and passed through a 0.22 um 96-well filter plate. The filtered supernatant (100 uL) was added to medium containing mAbs to achieve a final mAb concentration of 1 µg/ml in a total volume of 1 ml. The virus:antibody mixture was incubated for 1 hour at 37°C and used to inoculate 2×10^5^ HEK-293T/ACE2cl.22 cells for a second passage (p2) in the presence of antibodies. Medium was again replaced with fresh antibody-containing medium after 24 h, and the putatively selected p2 virus population was harvested after 48h. RNA was then extracted from 100 uL of filtered p2 supernatant and reverse transcribed using the SuperScript VILO cDNA Synthesis Kit (Thermo Fisher Scientific). Sequences encoding the extracellular domain of the spike protein were initially amplified using KOD Xtreme Hot Start Polymerase. In a subsequent PCR reaction, RBD-specific primers were used to amplify the receptor-binding domain (RBD), and the resulting products were purified and sequenced.

### Receptor binding domain sequencing

PCR products encoding the RBD were subjected to tagmentation reactions, that included 0.25 μl of Nextera TDE1 Tagment enzyme, 1.25 μl of TD Tagment buffer, and 10 ng of RBD PCR product. The tagmentation reaction was performed at 55ºC for 5 minutes. Next, 3.5 μl of KAPA HiFi HotStart ReadyMix and 1.25 μl of i5/i7 barcoded primers were added to the tagmented DNA to incorporate a unique indexing sequence. Following purification with AmpureXP beads, the barcoded DNA from several reactions was pooled together and denatured with 0.2 N NaOH. The barcoded library was then diluted to 12 pM and sequenced using Illumina MiSeq Nano 300 V2 cycle kits. Illumina sequencing reads were trimmed using the BBDuk function of Geneious Prime to remove adapter sequences and low-quality reads. The forward and reverse sequencing reads of each reaction were then aligned to the corresponding reference sequence and annotated for the presence of mutations. To detect RBD escape mutations, the minimum variant frequency (i.e.-the minimum fraction of reads that contained a substitution for it to be denoted as such) was set at 10%.

### Mutant spike expression plasmids and pseudotyped virus generation

Selected point mutations that arose in the rVSV/SARS-CoV-2 selection experiments were introduced into pCR3.1-based SARS-CoV-2 spike expression plasmids using a primer-based overlap extension approach as described above. Mutations that were selected in rVSV/SARS-CoV-2/GFP_2E1_ populations were introduced into the corresponding pCR3.1-based Wuhan-Hu-1 spike expression plasmid. Conversely, mutations that were selected in rVSV/SARS-CoV-2/GFP_BA1.1_, or rVSV/SARS-CoV-2/GFP_BA2_, were introduced in the context of all three (Wuhan-Hu-1 BA.1, BA.2) spike expression plasmids.

The generation of (HIV/NanoLuc) SARS-CoV-2 pseudotyped particles was described previously[24]. In brief, HEK-293T seeded in 10 cm dishes were transfected with 2.5 µg of a pCR3.1-SARS-CoV-2 spike expression plasmid and 7.5 µg of the pHIV-1_NL4-3_ ΔEnv-NanoLuc reporter virus plasmid using 44 µl polyethylenimine (PEI, 1 mg/ml) in 500 ml serum-free DMEM medium. The medium was changed 24 h post transfection. The virus containing supernatant was harvested at 48 h post transfection, passed through a 0.22 µm pore-size polyvinylidene fluoride syringe filter filter, aliquoted and stored at -80ºC. The titers were assessed on HT1080/ACE2cl.14 cells seeded 10^4^ cells/well in 100 µl medium in black flat-bottom 96-well plates. A 5-fold serial dilution was performed and 100 µl were transferred to the target plate. The plates were incubated for 48 h and NanoLuc luciferase activity determined as described below.

### Neutralization Assays

To assess the neutralization of pseudotyped HIV-1 viruses encoding the RBD point mutations that we had identified during our selections experiments by mAbs, neutralization assays were performed. First, HT1080 cells were seeded at 10^4^ cells/well in 100 µl of medium. The next day, 60 µl of diluted virus (adjusted to give ∼10^7^ relative light units in the infection assay) were incubated with 60 µl of mAb solution for 1 hour at 37°C in a 96-well plate. Each mutant virus was initially screened for neutralization by the entire panel of monoclonal antibodies at a single antibody concentration of 1 µg/ml infection relative to no-antibody controls determined. For viruses deemed to be resistant to antibody neutralization, defined as having a relative infection value >0.4 relative to the uninhibited virus control, IC_50_ values were determined for parental virus and escape mutant. In this case, five-fold dilutions of monoclonal antibody, starting at an initial concentration of 1 µg/ml, were incubated with the pseudotyped viruses. Thereafter, 100 µl of each antibody/virus mixture were added to the HT1080 cells and incubated at 37°C for 48 hours. NanoLuc luciferase activity was then determined.

### Reporter gene assays, curve fitting, and statistics

For NanoLuc luciferase assays, cells were washed twice with PBS and lysed with 30 µl of Cell Culture Lysis reagent. NanoLuc luciferase activity was determined using the NanoGlo Luciferase Assay System. Substrate in NanoGlo buffer (30 µl) was added to the cell lysate.

Luciferase activity was measured using a CLARIOstar Plus plate reader using 0.1s integration time. The RLU readings were expressed as a decimal fraction of those derived from cells infected with pseudovirus in absence of antibodies.

To determine titers of rVSV/SARS-CoV-2/GFP, infected cells were trypsinized, fixed with 2% paraformaldehyde, resuspended in PBS/F-68 and the percentage of GFP-positive cells determined using an Attune NxT flow cytometer equipped with a 96-well CytKick Max autosampler.

Data was analyzed using Prism V9.41 (GraphPad). NT_50_ and IC_50_ values were determined using four-parameter nonlinear regression curve fit to infectivity data measured as RLUs. The bottom values were set to zero, the top values to one. Statistical significance between groups was assessed by a two-tailed t-test using Welch’s correction to account for unequal variances.

---

**Supplementary Figure 1.**
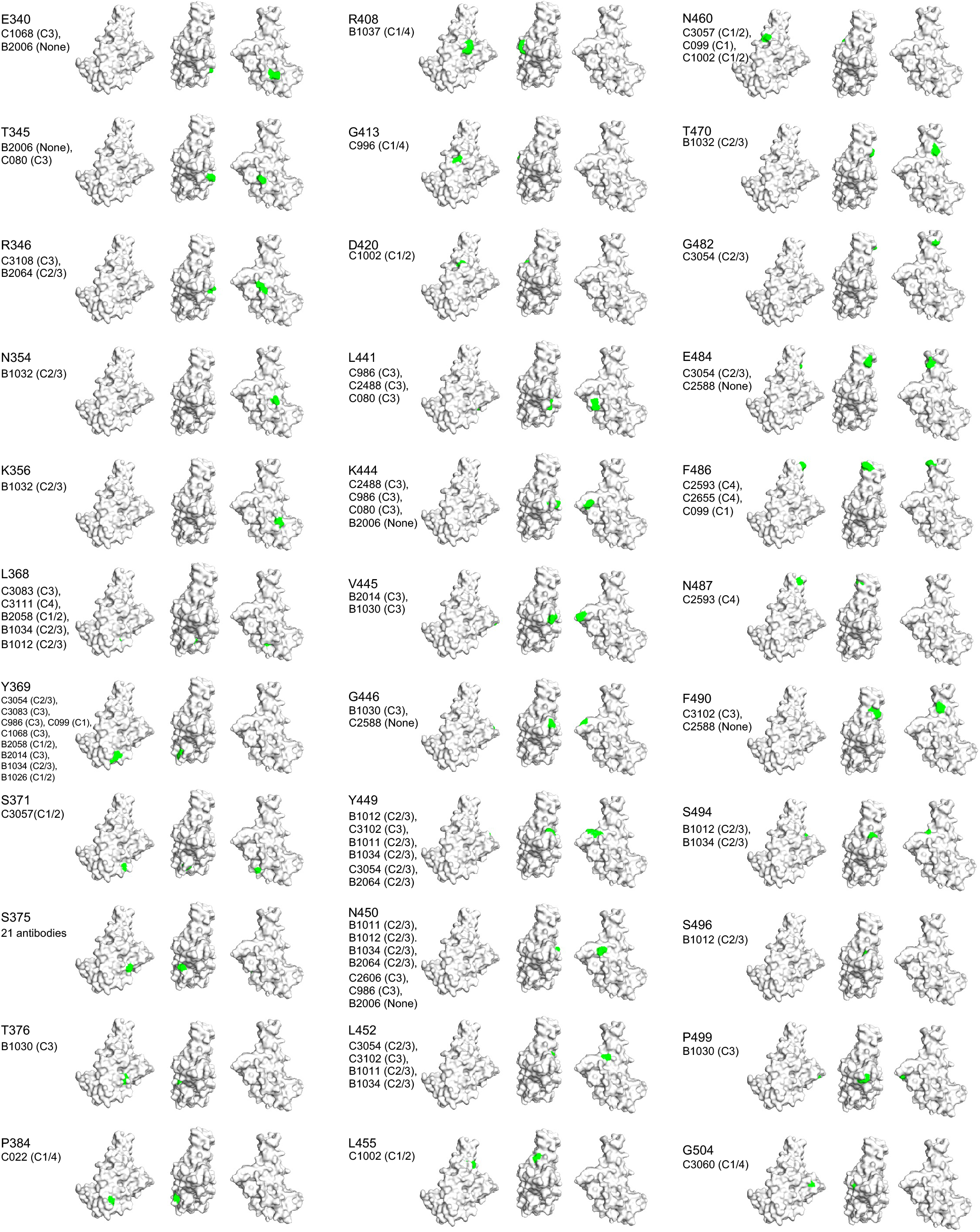
Substitutions enriched in rVSV/SARS-CoV-2 populations selected by broadly neutralizing antibodies. **a**, RBD structure (PDB ID : 7C8J) with positions (highlighted in green), at which substitutions occurring at frequencies of >10% were found after two passages of rVSV/SARS-CoV-2 encoding Wuhan-Hu-1, BA.1, and BA.2 spike proteins in the presence of 1 μg/ml of the indicated broadly neutralizing antibody whose class 1-4 designation is indicated.

**Supplementary Figure 2.**
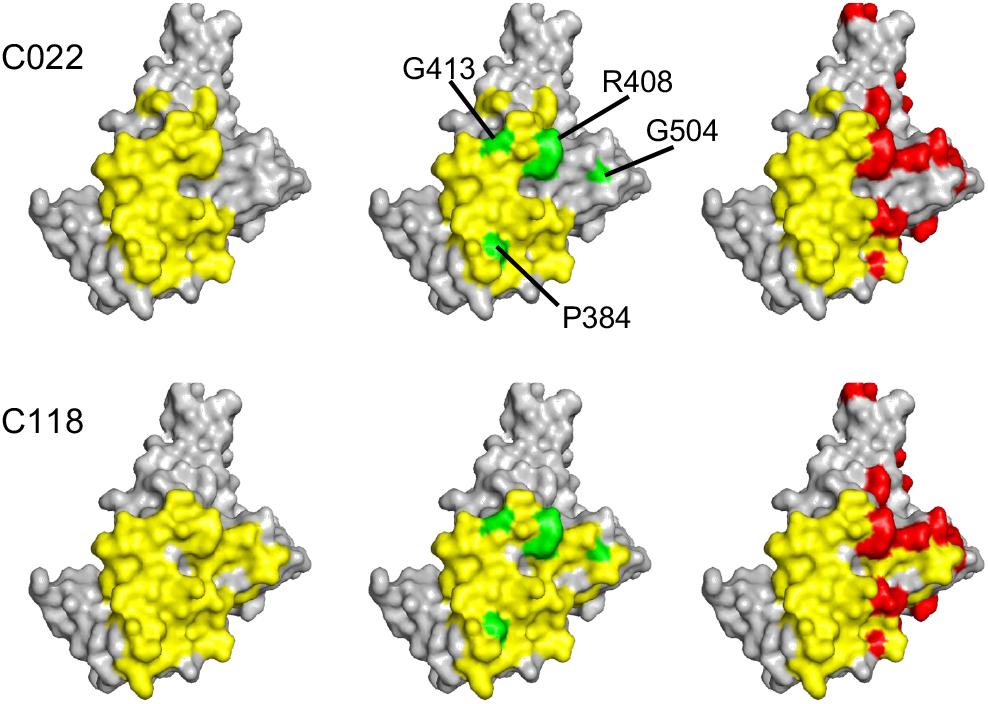
Resistance to broadly neutralizing class 4 and 1/4 antibodies. RBD structure (PDB ID : 7C8J) illustrating epitopes of two prototype class 4 antibodies (C022 and C118, yellow), substitutions that confer resistance to class 4 and 1/4 antibodies (green), and preexisting substitutions in BA.2 (red)

**Supplementary Figure 3.**
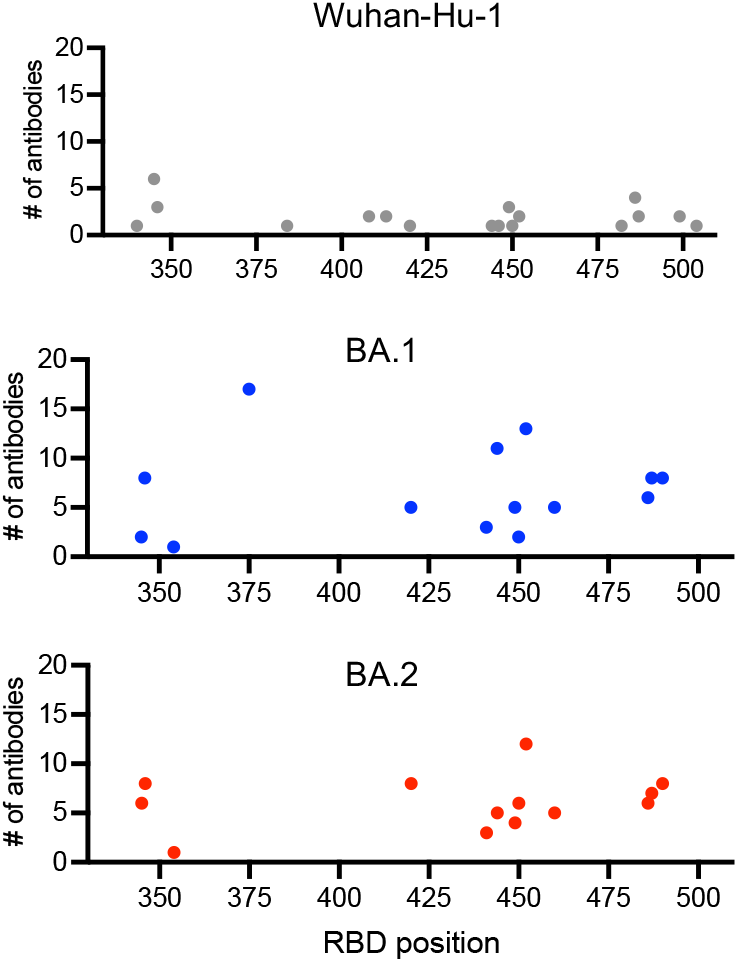
Context dependent effects of RBD substitutions on antibody escape. Number of broadly neutralizing antibodies for which substitutions in Wuhan-Hu-1, BA.1, and BA.2 backgrounds at positions along the length of the RBD confer escape. Antibody escape was defined as >5-fold increase in mutant pseudotype relative infection compared to parental pseudotype in the presence of 1μg/ml antibody and >10% relative infection compared to no antibody.

**Supplementary Figure 4.**
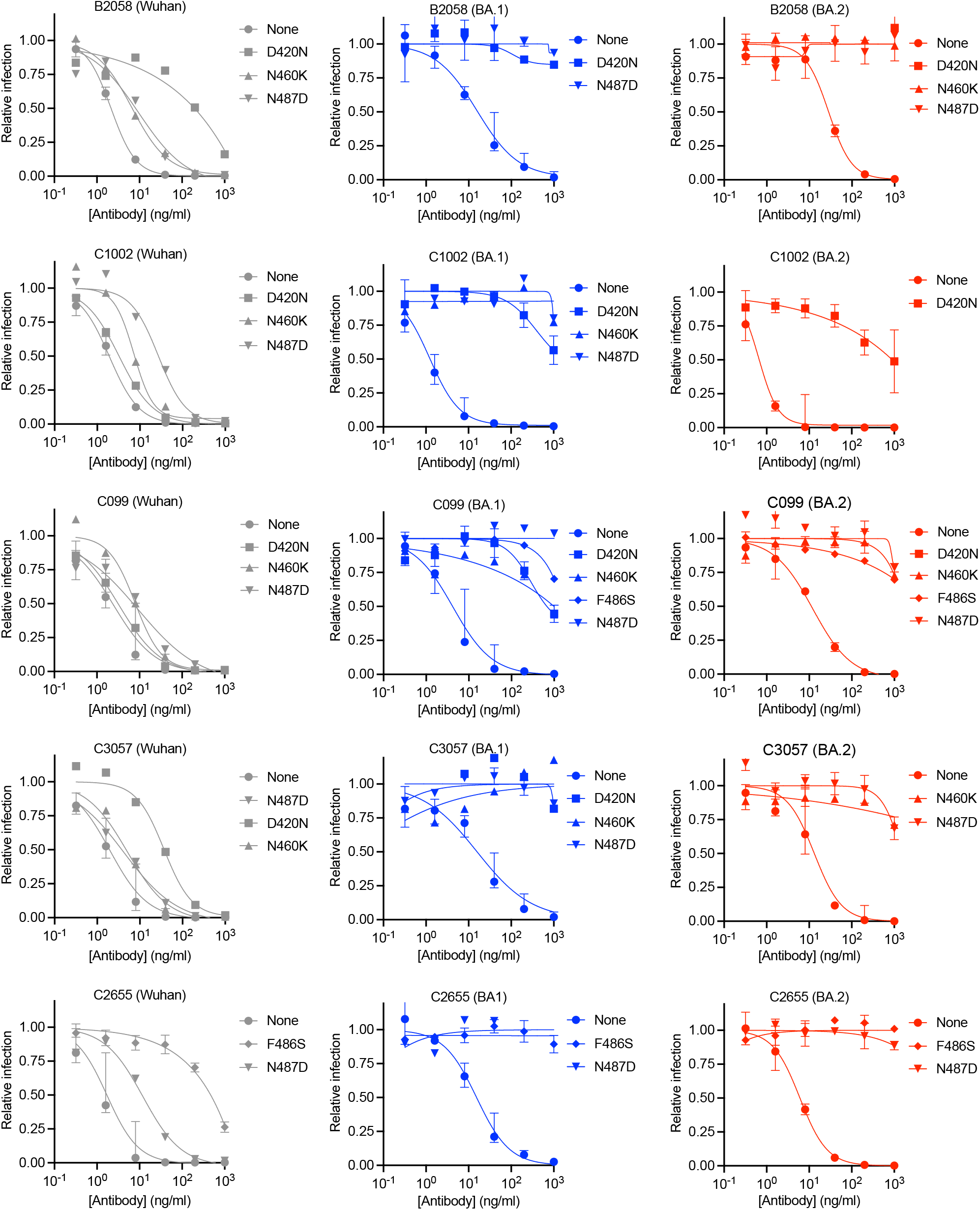
Epistatic effects of substitutions on broadly neutralizing class 1, 1/2 and 1/4 antibodies. Neutralization of RBD point mutant pseudotypes in Wuhan-Hu-1, BA.1, and BA.2 backgrounds by class 1, 1/2 and 1/4 antibodies. Median and range of 2 or 3 independent experiments is shown

**Supplementary 5.**
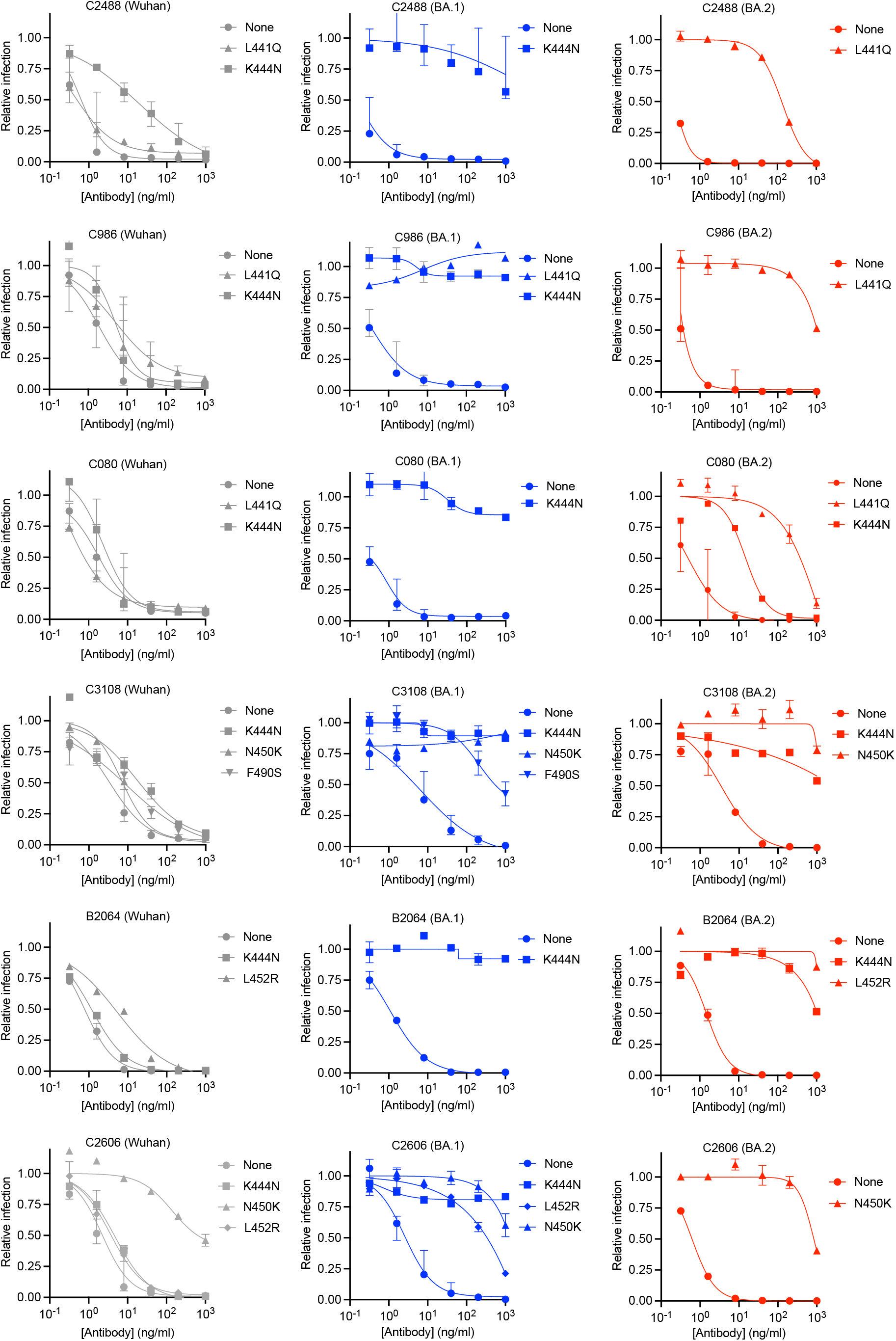
Epistatic effects of substitutions on broadly neutralizing class 2/3 and 3 antibodies (I) Neutralization of RBD point mutant pseudotypes in Wuhan-Hu-1, BA.1, and BA.2 backgrounds by class 2/3 and 3 antibodies. Median and range of 2 or 3 independent experiments is shown

**Supplementary 6.**
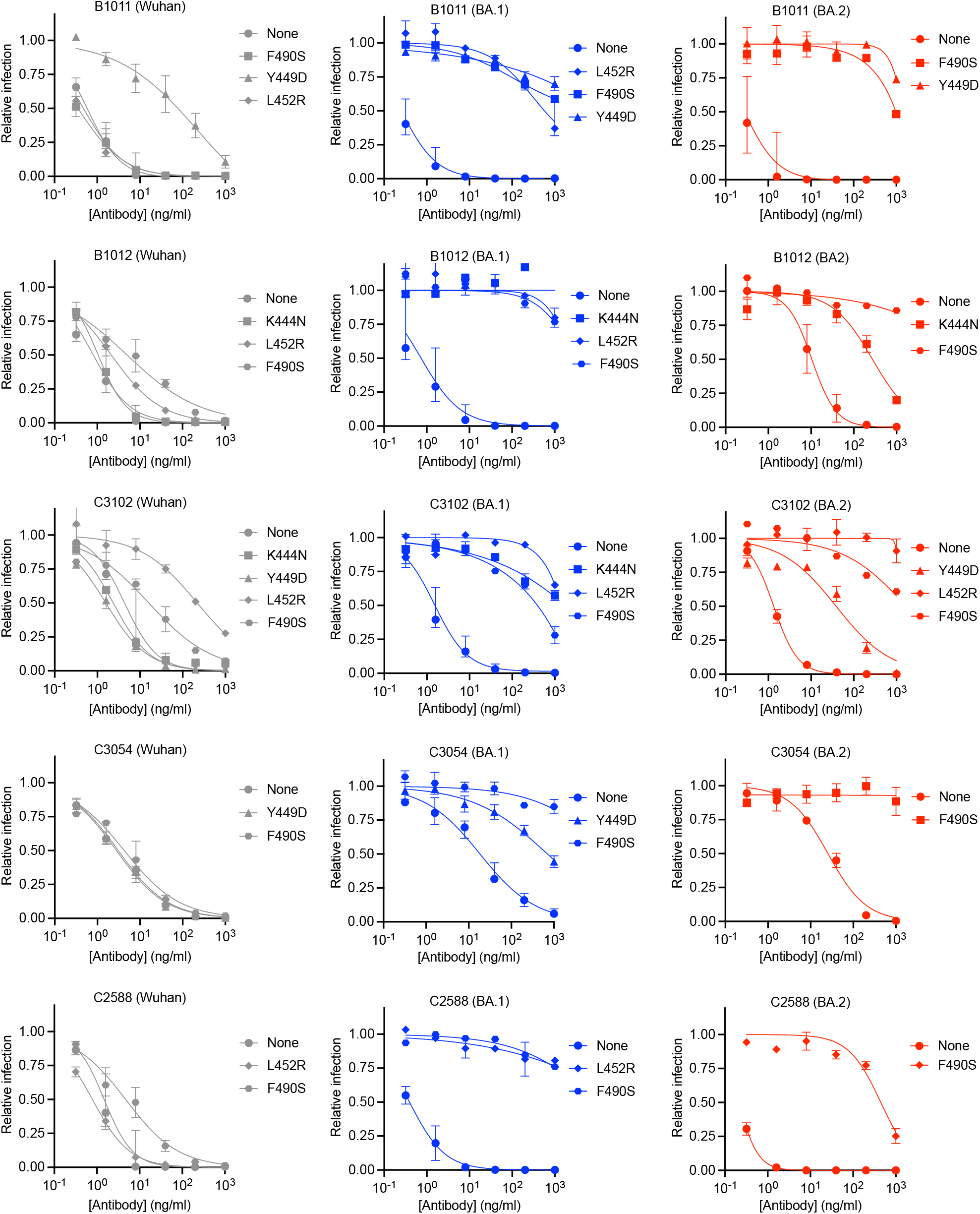
Epistatic effects of substitutions on broadly neutralizing class 2/3 and 3 antibodies (II) Neutralization of RBD point mutant pseudotypes in Wuhan-Hu-1, BA.1, and BA.2 backgrounds by class 2/3 and 3 antibodies. Median and range of 2 or 3 independent experiments is shown

**Supplementary Figure 7.**
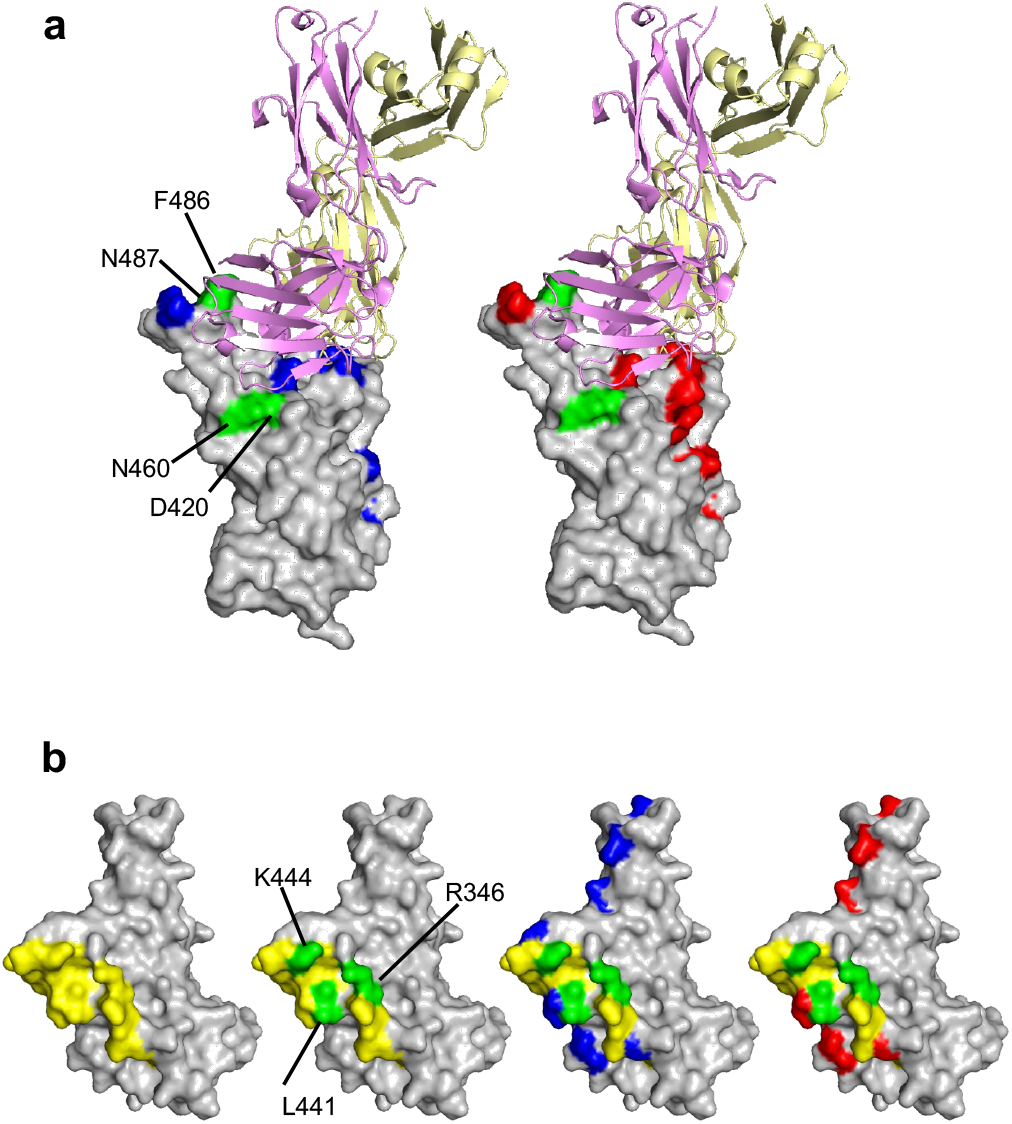
Pre-existing and escape substitutions in broadly neutralizing class 1 and class 3 antibody epitopes. **a)** Substitutions that confer context-dependent escape from the class I antibody C099, depicted on the C099:RBD complex structure (PDB ID 7R8L). Green indicates C099 escape substitutions. Blue and red indicate BA.1 and BA.2 substitutions, respectively. C099 heavy and light chains are magenta and yellow respectively. **b)** Substitutions that confer context-dependent escape from C080 class 3 antibody, depicted on RBD complex structure (PDB ID 7C8J). Yellow indicates the C032 (clonal ancestor of C080) epitope. Green indicates escape substitutions. Blue and red indicate BA.1 and BA.2 substitutions, respectively.

**Supplementary Figure 8.**
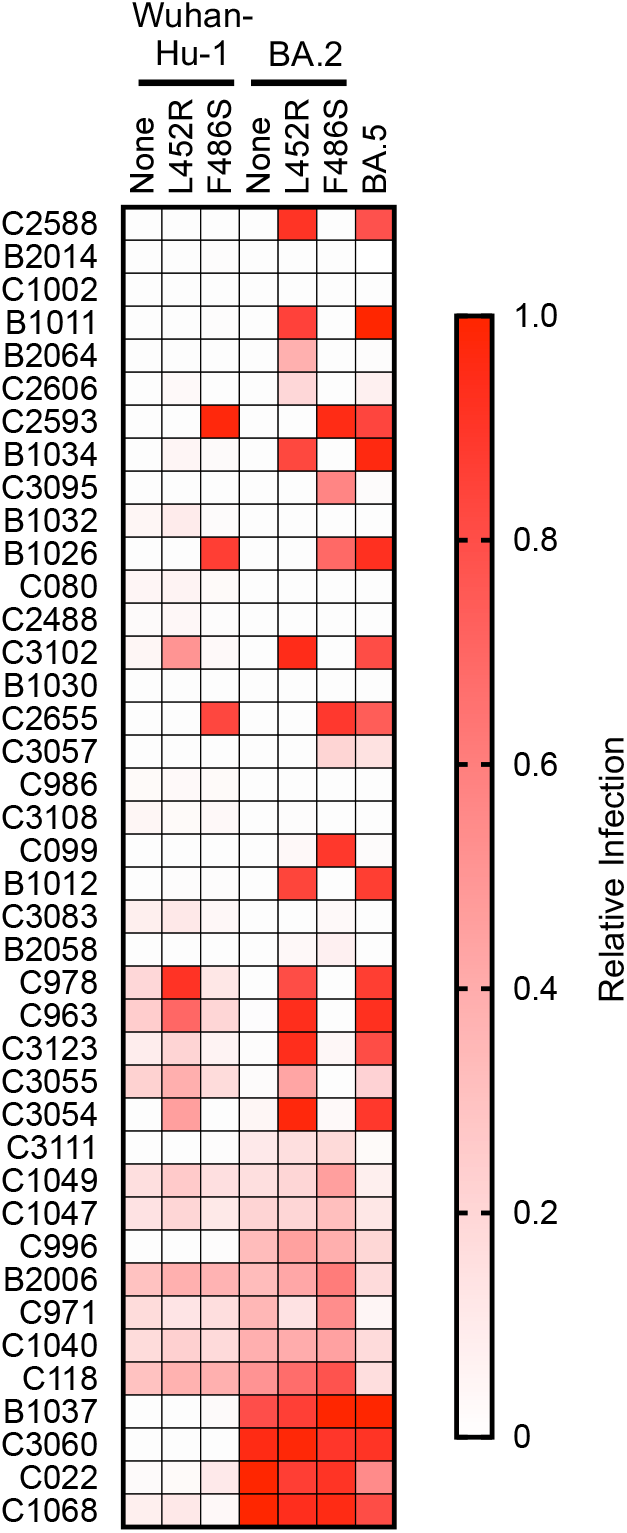
Context dependent effect of BA.5 substitutions (L452 and F486) on broadly neutralizing antibody sensitivity. Inhibition of Wuhan-Hu-1 and BA.2 RBD pseudotypes with L452R or F486S substitutions by broadly neutralizing antibodies, in comparison with BA.5. Relative infection is defined as the decimal fraction of infection measured (with 1 μg/ml antibody), relative to an uninhibited virus control (without antibody). Antibodies are listed in descending order of potency against BA.2 and median values from two independent experiments are displayed.

**Supplementary Data 1:** Demographics and SARS-CoV-2 clinical histories of participants

**Supplementary Data 2:** Properties and sequences of broadly neutralizing monoclonal antibodies

**Supplementary Data 3:** Frequencies of RBD substitutions identified during antibody selection experiments

**Supplementary Data 4:** Neutralization of Wuhan-Hu-1, BA.1, and BA.2 RBD point mutant pseudotypes by broadly neutralizing antibodies

**Supplementary Data 5:** p-values for Figure 4

## References

1. Weisblum Y, Schmidt F, Zhang F, DaSilva J, Poston D, Lorenzi JC, et al. Escape from neutralizing antibodies by SARS-CoV-2 spike protein variants. Elife. 2020;9. Epub 20201028. doi: 10.7554/eLife.61312. PubMed PMID: 33112236; PubMed Central PMCID: PMCPMC7723407.

2. Wibmer CK, Ayres F, Hermanus T, Madzivhandila M, Kgagudi P, Oosthuysen B, et al. SARS-CoV-2 501Y.V2 escapes neutralization by South African COVID-19 donor plasma. Nat Med. 2021;27(4):622–5. Epub 20210302. doi: 10.1038/s41591-021-01285-x. PubMed PMID: 33654292.

3. Planas D, Saunders N, Maes P, Guivel-Benhassine F, Planchais C, Buchrieser J, et al. Considerable escape of SARS-CoV-2 Omicron to antibody neutralization. Nature. 2022;602(7898):671–5. Epub 20211223. doi: 10.1038/s41586-021-04389-z. PubMed PMID: 35016199.

4. Schmidt F, Muecksch F, Weisblum Y, Da Silva J, Bednarski E, Cho AL, et al. Plasma Neutralization of the SARS-CoV-2 Omicron Variant. New Engl J Med. 2022;386(6):599–601. doi: 10.1056/NEJMc2119641. PubMed PMID: WOS:000736224400001.

5. Cao Y, Wang J, Jian F, Xiao T, Song W, Yisimayi A, et al. Omicron escapes the majority of existing SARS-CoV-2 neutralizing antibodies. Nature. 2022;602(7898):657–63. Epub 20211223. doi: 10.1038/s41586-021-04385-3. PubMed PMID: 35016194; PubMed Central PMCID: PMCPMC8866119.

6. Iketani S, Liu L, Guo Y, Liu L, Chan JF, Huang Y, et al. Antibody evasion properties of SARS-CoV-2 Omicron sublineages. Nature. 2022;604(7906):553–6. Epub 20220303. doi: 10.1038/s41586-022-04594-4. PubMed PMID: 35240676; PubMed Central PMCID: PMCPMC9021018.

7. Cele S, Jackson L, Khoury DS, Khan K, Moyo-Gwete T, Tegally H, et al. Omicron extensively but incompletely escapes Pfizer BNT162b2 neutralization. Nature. 2022;602(7898):654–6. Epub 20211223. doi: 10.1038/s41586-021-04387-1. PubMed PMID: 35016196; PubMed Central PMCID: PMCPMC8866126.

8. Zhou H, Dcosta BM, Landau NR, Tada T. Resistance of SARS-CoV-2 Omicron BA.1 and BA.2 Variants to Vaccine-Elicited Sera and Therapeutic Monoclonal Antibodies. Viruses. 2022;14(6). Epub 20220618. doi: 10.3390/v14061334. PubMed PMID: 35746806; PubMed Central PMCID: PMCPMC9228817.

9. Gaebler C, Wang Z, Lorenzi JCC, Muecksch F, Finkin S, Tokuyama M, et al. Evolution of antibody immunity to SARS-CoV-2. Nature. 2021;591(7851):639–44. Epub 20210118. doi: 10.1038/s41586-021-03207-w. PubMed PMID: 33461210; PubMed Central PMCID: PMCPMC8221082.

10. Cho A, Muecksch F, Schaefer-Babajew D, Wang Z, Finkin S, Gaebler C, et al. Anti-SARS-CoV-2 receptor-binding domain antibody evolution after mRNA vaccination. Nature. 2021;600(7889):517–22. Epub 20211007. doi: 10.1038/s41586-021-04060-7. PubMed PMID: 34619745; PubMed Central PMCID: PMCPMC8674133.

11. Turner JS, O’Halloran JA, Kalaidina E, Kim W, Schmitz AJ, Zhou JQ, et al. SARS-CoV-2 mRNA vaccines induce persistent human germinal centre responses. Nature. 2021;596(7870):109–13. Epub 20210628. doi: 10.1038/s41586-021-03738-2. PubMed PMID: 34182569; PubMed Central PMCID: PMCPMC8935394.

12. Muecksch F, Weisblum Y, Barnes CO, Schmidt F, Schaefer-Babajew D, Wang Z, et al. Affinity maturation of SARS-CoV-2 neutralizing antibodies confers potency, breadth, and resilience to viral escape mutations. Immunity. 2021;54(8):1853-68.e7. Epub 20210730. doi: 10.1016/j.immuni.2021.07.008. PubMed PMID: 34331873; PubMed Central PMCID: PMCPMC8323339.

13. Wang Z, Muecksch F, Schaefer-Babajew D, Finkin S, Viant C, Gaebler C, et al. Naturally enhanced neutralizing breadth against SARS-CoV-2 one year after infection. Nature. 2021;595(7867):426–31. Epub 20210614. doi: 10.1038/s41586-021-03696-9. PubMed PMID: 34126625; PubMed Central PMCID: PMCPMC8277577.

14. Muecksch F, Wang Z, Cho A, Gaebler C, Ben Tanfous T, DaSilva J, et al. Increased memory B cell potency and breadth after a SARS-CoV-2 mRNA boost. Nature. 2022;607(7917):128–34. Epub 20220421. doi: 10.1038/s41586-022-04778-y. PubMed PMID: 35447027; PubMed Central PMCID: PMCPMC9259484.

15. Greaney AJ, Loes AN, Crawford KHD, Starr TN, Malone KD, Chu HY, et al. Comprehensive mapping of mutations in the SARS-CoV-2 receptor-binding domain that affect recognition by polyclonal human plasma antibodies. Cell Host Microbe. 2021;29(3):463-76.e6. Epub 20210208. doi: 10.1016/j.chom.2021.02.003. PubMed PMID: 33592168; PubMed Central PMCID: PMCPMC7869748.

16. Schmidt F, Weisblum Y, Rutkowska M, Poston D, DaSilva J, Zhang F, et al. High genetic barrier to SARS-CoV-2 polyclonal neutralizing antibody escape. Nature. 2021;600(7889):512–6. Epub 20210920. doi: 10.1038/s41586-021-04005-0. PubMed PMID: 34544114; PubMed Central PMCID: PMCPMC9241107.

17. Barnes CO, Jette CA, Abernathy ME, Dam KA, Esswein SR, Gristick HB, et al. SARS-CoV-2 neutralizing antibody structures inform therapeutic strategies. Nature. 2020;588(7839):682–7. Epub 20201012. doi: 10.1038/s41586-020-2852-1. PubMed PMID: 33045718; PubMed Central PMCID: PMCPMC8092461.

18. Jette CA, Cohen AA, Gnanapragasam PNP, Muecksch F, Lee YE, Huey-Tubman KE, et al. Broad cross-reactivity across sarbecoviruses exhibited by a subset of COVID-19 donor-derived neutralizing antibodies. Cell Rep. 2021;36(13):109760. Epub 20210908. doi: 10.1016/j.celrep.2021.109760. PubMed PMID: 34534459; PubMed Central PMCID: PMCPMC8423902.

19. Huang M, Wu L, Zheng A, Xie Y, He Q, Rong X, et al. Atlas of currently available human neutralizing antibodies against SARS-CoV-2 and escape by Omicron sub-variants BA.1/BA.1.1/BA.2/BA.3. Immunity. 2022;55(8):1501-14.e3. Epub 20220615. doi: 10.1016/j.immuni.2022.06.005. PubMed PMID: 35777362; PubMed Central PMCID: PMCPMC9197780.

20. Cameroni E, Bowen JE, Rosen LE, Saliba C, Zepeda SK, Culap K, et al. Broadly neutralizing antibodies overcome SARS-CoV-2 Omicron antigenic shift. Nature. 2022;602(7898):664–70. Epub 20211223. doi: 10.1038/s41586-021-04386-2. PubMed PMID: 35016195.

21. Gaebler C, DaSilva J, Bednarski E, Muecksch F, Schmidt F, Weisblum Y, et al. Severe Acute Respiratory Syndrome Coronavirus 2 Neutralization After Messenger RNA Vaccination and Variant Breakthrough Infection. Open Forum Infect Dis. 2022;9(7):ofac227. Epub 20220507. doi: 10.1093/ofid/ofac227. PubMed PMID: 35818364; PubMed Central PMCID: PMCPMC9129198.

22. Wang Z, Zhou P, Muecksch F, Cho A, Tanfous TB, Canis M, et al. Memory B cell responses to Omicron subvariants after SARS-CoV-2 mRNA breakthrough infection. BioRxiv. 2022. doi: https://doi.org/10.1101/2022.08.11.503601.

23. Robbiani DF, Gaebler C, Muecksch F, Lorenzi JCC, Wang Z, Cho A, et al. Convergent antibody responses to SARS-CoV-2 in convalescent individuals. Nature. 2020;584(7821):437–42. Epub 20200618. doi: 10.1038/s41586-020-2456-9. PubMed PMID: 32555388; PubMed Central PMCID: PMCPMC7442695.

24. Schmidt F, Weisblum Y, Muecksch F, Hoffmann HH, Michailidis E, Lorenzi JCC, et al. Measuring SARS-CoV-2 neutralizing antibody activity using pseudotyped and chimeric viruses. J Exp Med. 2020;217(11). doi: ARTN e20201181 10.1084/jem.20201181. PubMed PMID: WOS:000587422000019.

25. Tegally H, Moir M, Everatt J, Giovanetti M, Scheepers C, Wilkinson E, et al. Emergence of SARS-CoV-2 Omicron lineages BA.4 and BA.5 in South Africa. Nat Med. 2022. Epub 20220627. doi: 10.1038/s41591-022-01911-2. PubMed PMID: 35760080.

26. https://gisaid.org.

27. Viant C, Weymar GHJ, Escolano A, Chen S, Hartweger H, Cipolla M, et al. Antibody Affinity Shapes the Choice between Memory and Germinal Center B Cell Fates. Cell. 2020;183(5):1298-311.e11. Epub 20201029. doi: 10.1016/j.cell.2020.09.063. PubMed PMID: 33125897; PubMed Central PMCID: PMCPMC7722471.

28. Cao Y, Yisimayi A, Jian F, Song W, Xiao T, Wang L, et al. BA.2.12.1, BA.4 and BA.5 escape antibodies elicited by Omicron infection. Nature. 2022. Epub 20220617. doi: 10.1038/s41586-022-04980-y. PubMed PMID: 35714668.

29. Kaku CI, Bergeron AJ, Ahlm C, Normark J, Sakharkar M, Forsell MNE, et al. Recall of preexisting cross-reactive B cell memory after Omicron BA.1 breakthrough infection. Sci Immunol. 2022;7(73):eabq3511. Epub 20220729. doi: 10.1126/sciimmunol.abq3511. PubMed PMID: 35549299; PubMed Central PMCID: PMCPMC9097882.

30. Wang Z, Schmidt F, Weisblum Y, Muecksch F, Barnes CO, Finkin S, et al. mRNA vaccine-elicited antibodies to SARS-CoV-2 and circulating variants. Nature. 2021;592(7855):616–22. Epub 20210210. doi: 10.1038/s41586-021-03324-6. PubMed PMID: 33567448; PubMed Central PMCID: PMCPMC8503938.

